# Optimal Social Group Size in Spotted Hyaenas (*Crocuta crocuta*): Insights into a Multilevel Society

**DOI:** 10.1101/2023.09.19.558384

**Authors:** R.I.M. Dunbar

**Affiliations:** Department of Experimental Psychology University of Oxford Radcliffe Quarter Oxford OX2 6GG UK

**Keywords:** fractal structure, infertility trap, stress, females alliances, benefits of group-living

## Abstract

The spotted hyaena lives in unusually large social groups for a carnivore. Since the infertility trap normally limits the size of social groups in mammals, it seems likely that this species has evolved some way of mitigating the stresses involved. In primates, this usually takes the form of female-female alliances, often embedded in multilevel social systems. I show (1) that the distribution of hyaena clan sizes is multimodal, with a fractal scaling close to 3 and a base unit of 12-15 individuals (3-5 reproductive females) and (2) that fertility is a trade off between the benefits of having more males in the group and the costs incurred by having more females, with 4-5 as the limit on the number of females that can live together without their reproductive rates falling below the demographic replacement rate. I present evidence that females buffer themselves against the infertility trap by forming matrilineal alliances that in turn create a multilevel structure. In this respect, hyaena resemble cercopithecine primates in using social strategies to enable animals to live in larger groups than they would otherwise be able to do.

## Introduction

Mammal social groups characteristically take one of two forms: aggregations (temporary herds) or congregations (stable, bonded social groups). The first is characterised by a unimodal (normal or Poisson) distribution where the mean size is determined by the species’ typical habitat conditions; the second is characterised by a multimodal, fractal distribution formed by a series of Poisson distributions that partially overlie each other. The second arises because, unlike aggregations, bonded groups cannot easily lose members once their size exceeds the optimum; instead, they have to wait until the group is large enough to undergo fission so as to produce daughter groups of a minimum size to ensure survival (Dunbar et al. 2009). When habitat conditions demand larger groups, these are created by delaying fission, thereby creating groups that are a multiple of the base unit (Dunbar & Shultz 2021a).

While most mammal societies are of the first kind, primates are characterised by societies of the second kind (Shultz & Dunbar 2007, 2010). Not only are species mean group sizes fractally distributed (Dunbar et al. 2018a), but their groups also have a fractal internal structure (Hill et al. 2008; Dunbar 2020; Escribano et al. 2022). In primates, the fractal structure is created by a local trade off between an infertility trap to which all mammals are subject and the benefits of being in large groups, mediated by a capacity to form coalitions that buffer the females against the stresses that induce infertility (Dunbar & Shultz 2021b). While the fact that primate groups are fractally structured is well established (Dunbar et al. 2018a; Dunbar & Shultz 2021a), the case for similarly organised societies in other mammals is less clear, not least because most species with bonded social systems typically have small groups (equids, elephants, lion, tylopods, mustelids). However, evidence for fractal structuring has been adduced for elephants and orcas (Hill et al. 2008), although the basis for this remains unknown.

The spotted hyaena (*Crocuta crocuta*) provides a promising example of a primate-like social organisation. It lives in clans whose members are usually related to each other (Holecamp et al. 2012), and share a common hunting territory and set of dens (Boydston et al. 2006). Clans can, however, vary in size between about 5 and several hundred individuals (Holecamp & Dloniak 2010). Other than to suggest that larger groups allow hyaena to hunt more profitable, large-bodied prey (Kruuk 1972; Höner et al. 2002), few studies comment on the fact that hyaena clans vary so much in size (3-150 individuals: Holecamp & Dloniak 2010) both within and between habitats or ask whether there is an optimal (or characteristic) size for the species. Although prey abundance has been offered as an explanation for large clans, the fact that clan sizes can vary from less than a dozen to more than 70 within the same habitat suggests that other forces must be at work. In *Papio* baboons, whose groups vary across a similar size range, analysis of the distribution of group sizes suggests that this is multimodal, forming two distinct optima whose size is determined by the habitat level of predation risk (Dunbar et al. 2018b; Dunbar & MacCarron 2019). More importantly, mathematical analysis of the efficiency of information flow through networks suggests that only certain group sizes are optimal in this case, with these optima forming a distinct series (5, 15, 50, 150, 500) with a scaling ratio of ∼3 (West et al. 2020, 2023). In addition to defining the characteristic sizes of primate social groups (Dunbar et al. 2018b), these values also form the template for the internal structuring of large social groups (Dunbar 2020; Escribano et al. 2022).

To determine whether spotted hyaena have a formal multilevel social system like that found in anthropoid primates, I first determine whether or not the size distribution of hyaena clans forms a single unimodal (normal or Poisson) distribution. To determine whether the hyaenas suffer from the infertility trap, and whether this places a limit on clan size, I examine fertility rates as a function of group size. I then seek to establish whether hyaena use alliances to overcome the infertility trap in order to allow larger clans to be stable. Finally, since populations differ considerably in their characteristic clan sizes, I seek to determine why some populations might prefer to live in very large groups. The answers to these questions both help us to understand the nature of hyaena society, and offer additional insights into the processes that allow some species to live in large groups despite the infertility trap.

## Methods

I searched the literature for data on clan sizes in *Crocuta crocuta*. I located 20 studies, all of which had involved at least a year of field study of known individuals. In three cases, studies that were carried out on the same population at least two decades apart were treated as independent samples on the grounds that there was ample time for their population structure to change between successive samples. The clan sizes given by these studies are based either on census data at a specific time (usually the beginning or end of the study) or on the average over a specified period of time. The average number of clans censused at each site was 3.1±2.3, yielding a total of 57 clans with reliable census data.

To determine whether there is an optimal clan size, I plot the cumulative distribution of all 57 clan sizes and determine whether this is continuous (i.e. a sigmoid shape reflecting a normal distribution) or exhibits a ‘broken stick’ pattern (involving a distinct change in slope) indicating that the data form two (or more, if there are several changes in slope) independent distributions (MacArthur 1957; Magurran 1988). A change in slope identifies the optimal group size (Dunbar & Shultz 2023). To determine the number and mean size of clusters, I use *k*-means cluster analysis, versions of which has been widely used to detect fractal layering in human and primate social networks (Arnaboldi et al. 2015; Kordsmeyer et al. 2017; Dunbar et al. 2021a,b). This seeks to partition the distribution into natural bins based on optimising gaps in the sequence of observed values. There are no formal solutions to determining the optimal number of clusters, but convention is to minimise the number of clusters that maximises fit (Coulson 1987). To identify this, I plot the goodness-of-fit (indexed by the F-value) for different numbers of clusters. We seek the cluster number where the fit ceases to increase significantly.

To determine whether hyaena experience an infertility trap, I plot fertility against group demographics in two separate samples. First, I extracted from the dataset all studies that listed the number of cubs as well as number of adult males and adult females for individual clans, and calculated mean cubs per adult female. In primates, fertility is a trade off between a positive benefit of total group size and a negative function of the number of females (the infertility trap); I test for a similar relationship in hyaenas with a multiple regression with mean fertility in a clan (indexed as cubs/female) as the outcome variable, and number of adult males, number of adult females and total clan size (including subadults) as predictors. I then use data given by Holecamp et al. (2012) for one large clan to compare lifetime reproductive output (estimated from long term pedigrees) for females of different dominance rank to determine whether (a) the infertility trap applies within as well as between clans and (b) whether partitioning females into matrilineal alliances alleviates the infertility trap. In both cases, only negative correlations are of interest, so a 1-tailed test of significance is appropriate.

To determine gain some insight into why some populations live in much larger clans than others, I ran a regression of mean clan size at different locations against climatic variables (mean ambient temperature and mean annual rainfall). Data on climate at individual sites are taken from either Holecamp & Dloniak (2011) or original sources.

All the data are given in *Supplementary Datafiles 1* (clan demography at different sites) and *2* (completed family size for individual females in the Talek clan from the Serengeti, given by Holecamp et al. 2012).

All statistical analyses were carried out using SSPS v.29.

## Results

Fig. 1 plots the frequency distribution of clan sizes, with a spline fit (solid line) and an overlying normal distribution (dashed line). Mean size is 33.1±25.9SD. A unimodal distribution is clearly a poor fit to the data (Kolmogorov-Smirnov one sample test: normal, p<0.001; Poisson, p<0.01; 2-tailed tests). The spline fit suggests a trimodal distribution. A *k*-means cluster analysis with 2≥*k*≥6 yields plausible solutions at *k*=2 and, especially, *k*=3, but models with more than three clusters do not add significantly to the goodness-of-fit (Fig. 2). A 2-cluster solution gives cluster means of 15.7 and 62.3 (with N=37 and 20 clans in each cluster), while the 3-cluster solution gives cluster means of 11.3, 41.7 and 71.8 (with N=30, 12 and 15 clans, respectively). In essence, adding additional clusters simply partitions the larger of the *k*=2 clusters, leaving the smaller cluster relatively untouched.

**Fig. 1.**
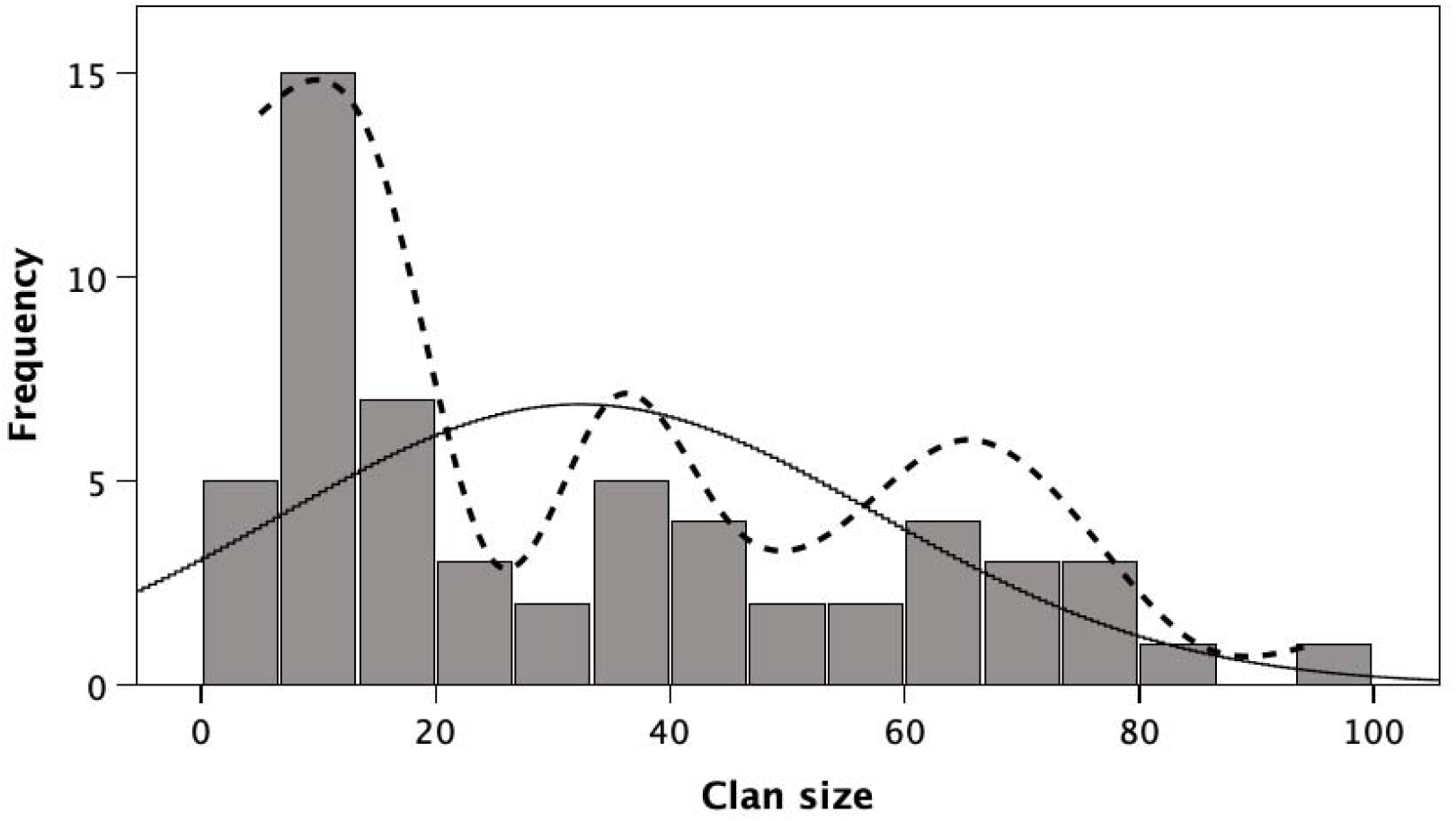
Distribution of spotted hyaena clan sizes, with fits for a normal distribution (solid line) and a third-order spline (dashed line).

**Fig. 2.**
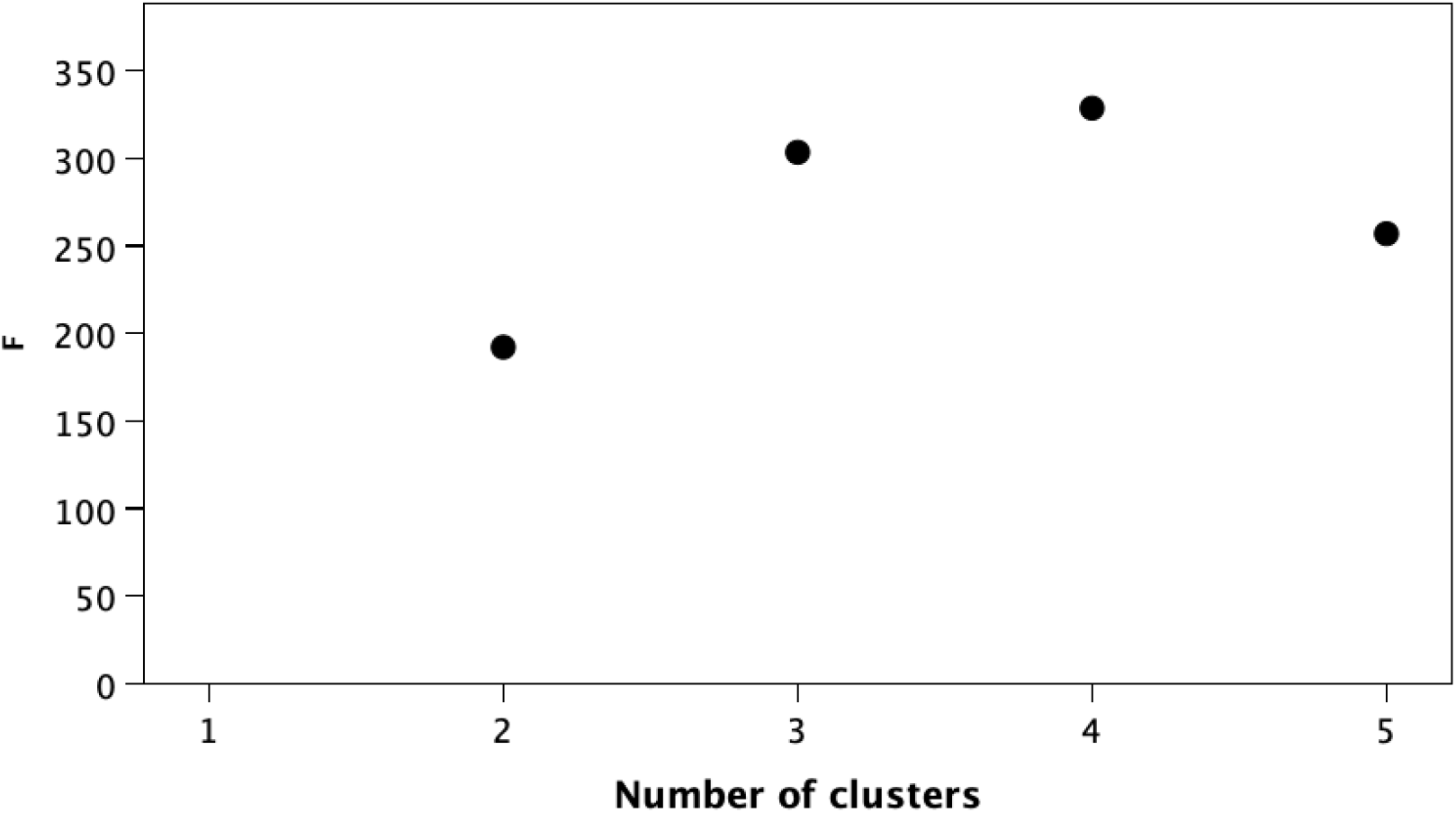
The optimal number of clusters in the distribution of hyaena clan sizes shown in Fig. 1 is indicated by the changing pattern in the goodness-of-fit (indexed as F-ratio) for a *k*-means cluster analysis with 2≤*k*≤5. There is minimal further improvement in goodness-of-fit after *k*=3.

In broad agreement with these results, a ‘broken stick’ solution yields a natural division between small and large clusters at a clan size of exactly 15 (Fig. 3). This value sets an upper limit on the size of the smaller cluster, since it defines the transition between two basic types of group (small and large). This suggests that spotted hyaena are characterised by a base social grouping (clan) of 12-15 adults and subadults, with larger groupings being created when desirable by deferring clan fission so as to allow two or more base units to live together as a mega-clan. The cluster analysis indicates a scaling ratio of λ=2.7, close to the λ=3 scaling ratio found in primate and human social networks.

**Fig. 3.**
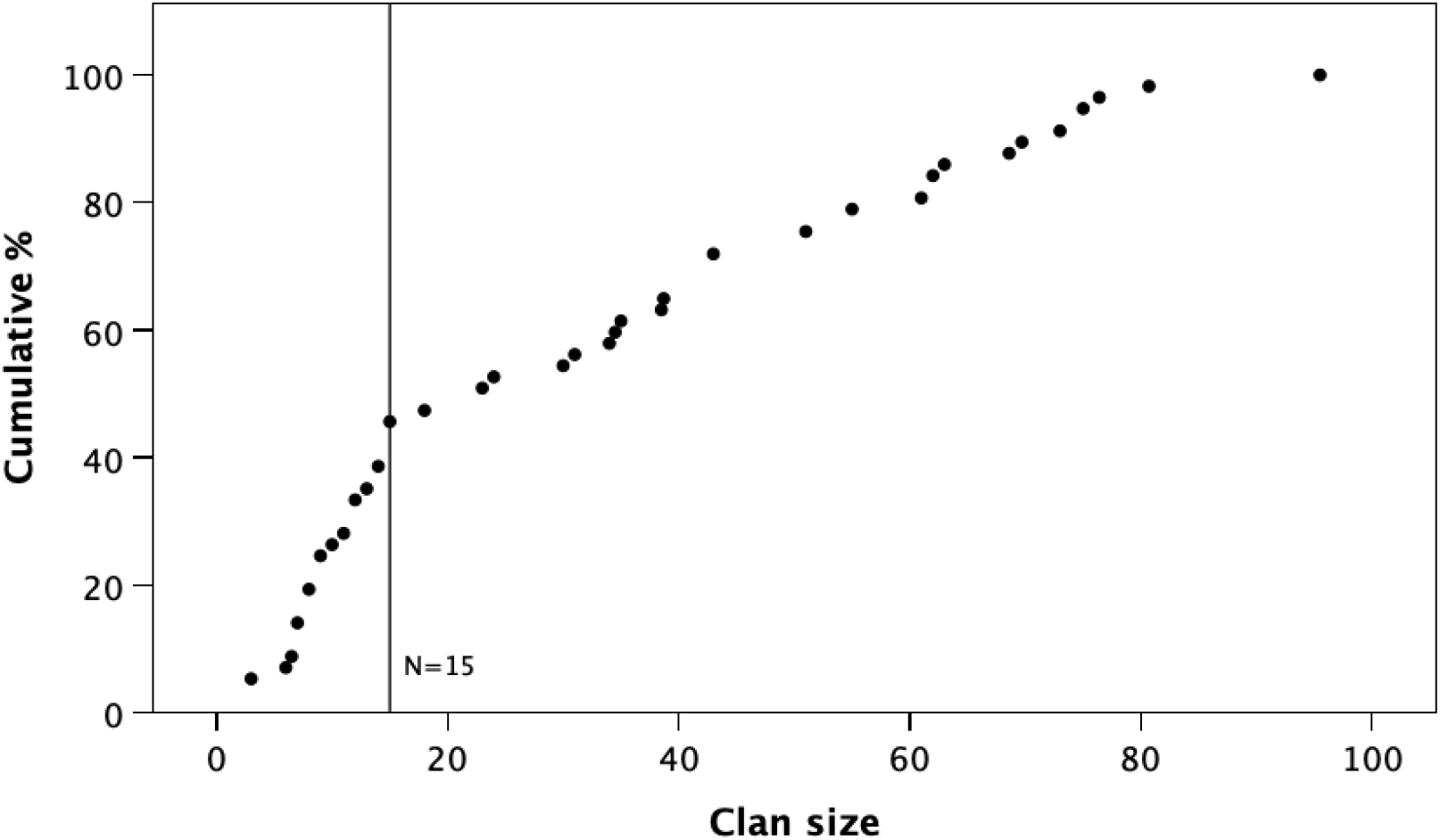
The classic ‘broken stick’ method applied to the cumulative distribution of hyaena clan sizes identifies a natural phase shift that partitions the distribution into two regions (see text for details).

To test whether this pattern is underpinned by an ‘infertility trap’, I first plotted mean birth rate per female against clan size (Fig. 4a) to determine whether this is a simple negative linear relationship (as in socially less complex mammals) or ∩-shaped (as in anthropoid primates and other mammals with bonded social systems) (see Dunbar & Shultz 2021b). Overall, there is no significant relationship (linear: r^2^=0.050, F_1,19_=0.99, p=0.331; quadratic: r^2^=0.106, F_2,18_=1.07, p=0.363). It is, however, obvious that the East African populations differ radically from the southern African populations. Separating these out yields a near-significant negative power relationship for the southern populations (r^2^=0.474, F_1,5_=4.51, p=0.087) and a significant positive power relationship in the East African populations (r^2^=0.494, F_1,12_=11.69, p=0.005), both indicating asymptotic relationships. It is possible that the East African populations are entering a downward trajectory on the right hand side of the graph that would, if there were data from clans >80 in size, yield a ∩-shaped graph. Notice that only one clan has a birth rate that falls below the demographic replacement rate.

**Fig. 4.**
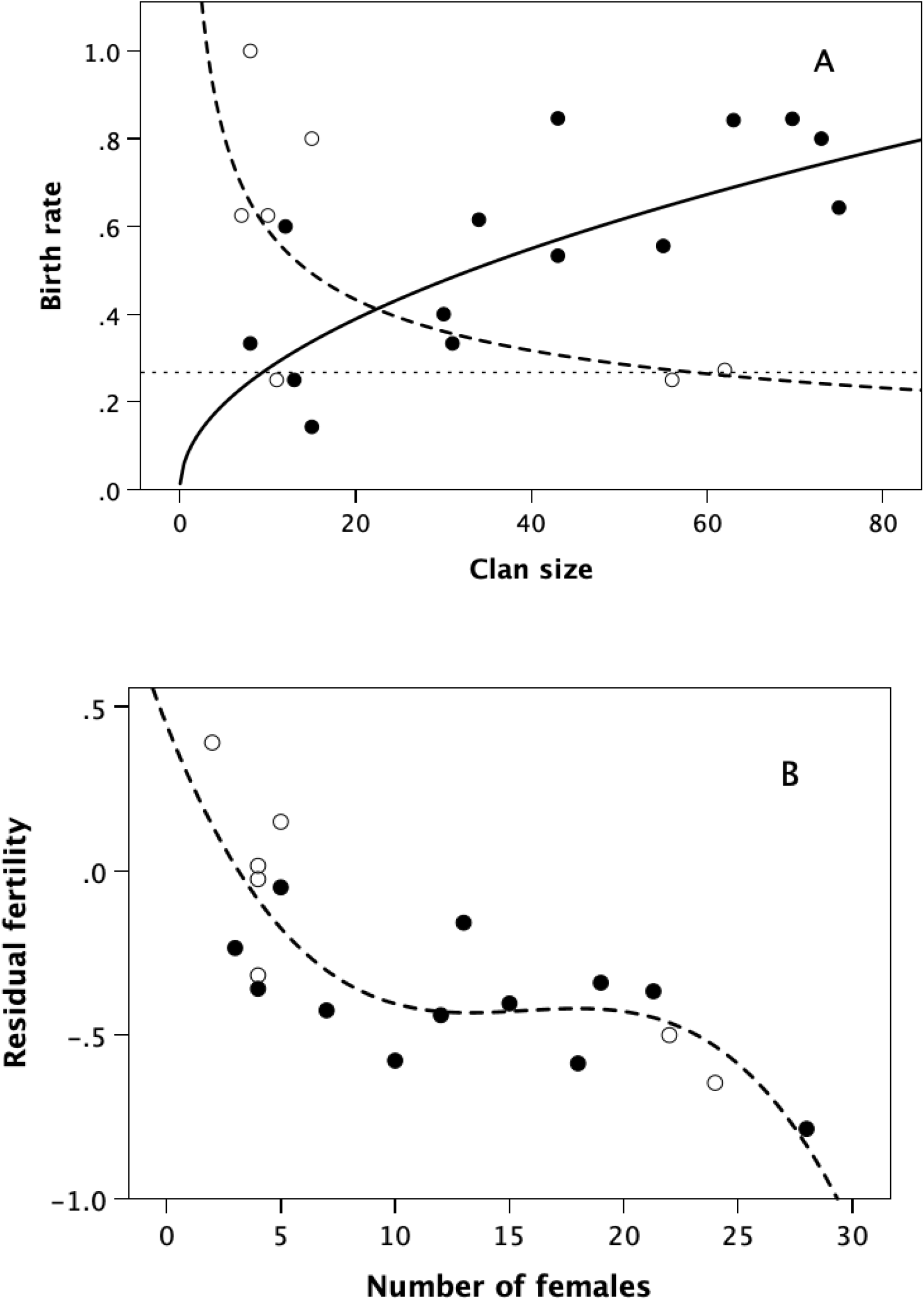
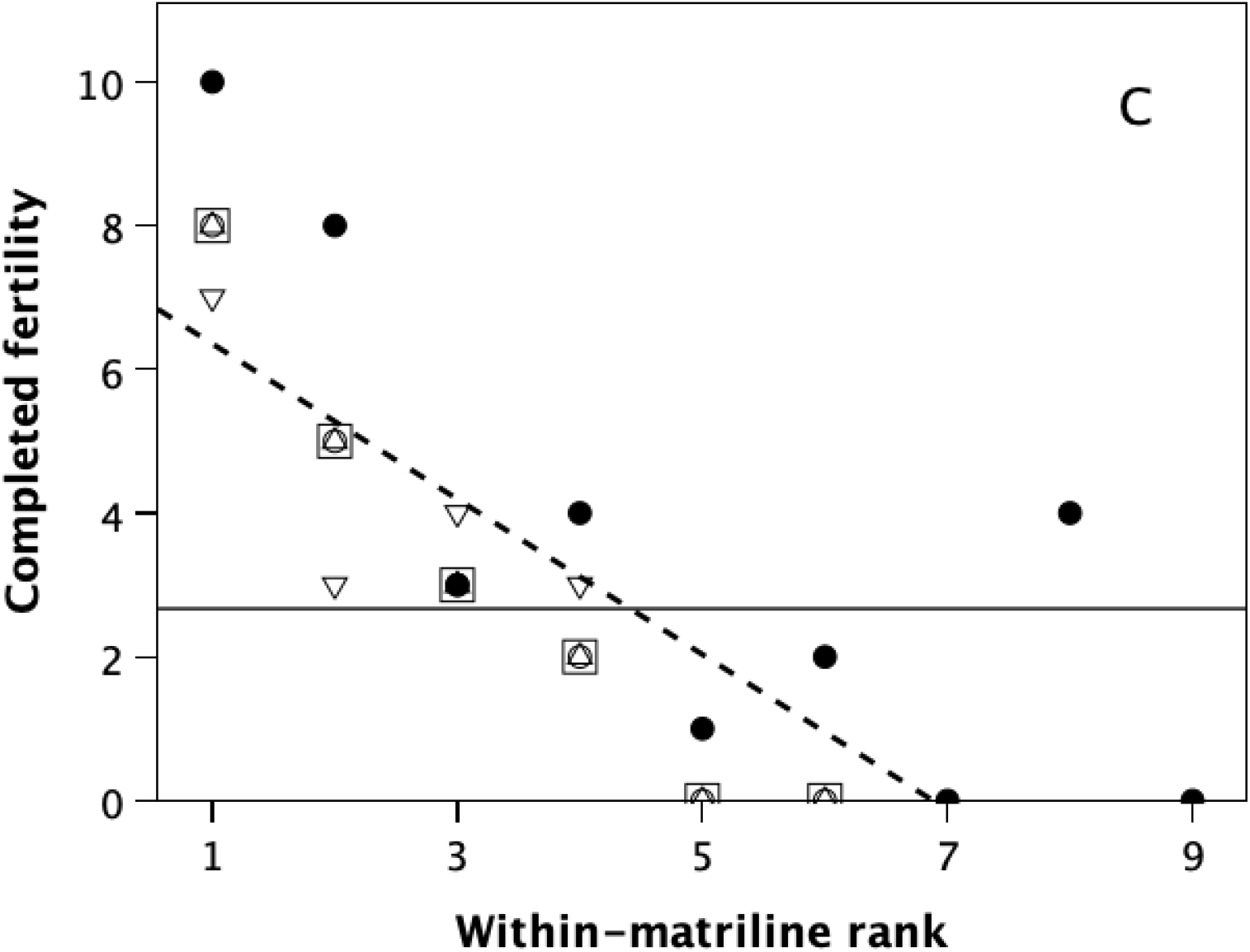
(a) Mean birth rate (estimated as the number of cubs per adult female, for cubs that emerged from the den at ∼4 weeks of age) plotted against clan size, for individual East African (filled symbols) and southern African (unfilled symbols) clans. The thin dotted line demarcates the demographic replacement rate: when mean birth rate is below this line, clans will decline in size. The replacement rate allows for cub mortality to 18 months of 25% (Hofer & East 2003). (b) Residual mean birth rate (adjusted for the number of adult males in the clan) as a function of the number of adult females for individual East African (filled symbols) and southern African (unfilled symbols) clans. The best fit regression is a cubic equation (r^2^=0.635, F_3,15_=8.70, p=0.001). (c) Number of cubs for individual females in the Serengeti Talek clan as a function of female dominance rank within their matriline. Matrilines are indicated by different symbols (solid circles: dominant matriline; unfilled squares: most subordinate matriline).

To check whether there might be an underlying ∩-shaped distribution created by a trade off between different demographic variables (the basis of the ∩-shaped distributions in primates and some other mammals), I ran a multiple regression analysis on the mean number of cubs per adult female in clans of different size and composition (Table 1). Model 1 included all three key demographic variables (number of adult males, number of adult females, and total clan size including subadults). Model 2 drops the least significant predictor (total clan size). Only Model 2 is statistically significant, and it is the only model for which all slope coefficients are individually significant. Model 2 suggests that fertility (indexed by the number of cubs that survive to emergence from the den at about four weeks of age) is positively related to the number of adult males in the clan, but negatively with the number of adult females. In other words, as in the case of primates, there is a benefit to be derived from living in larger clans (in this case, mainly due to the number of extra males), but a price to pay for doing so as a consequence of the increased number of females.

**Table 1.** Multiple regression models predicting fertility (indexed as census-dated cubs per adult female in 18 different clans)

| Model | Intercept | Slope coefficients |  | Total clan† |  | F | df | p |
| --- | --- | --- | --- | --- | --- | --- | --- | --- |
|  |  | Males | Females |  |  |  |  |  |
| 1 | 0.528 | 0.029 | -0.058 | 0.014 |  | 3.00 | 3,15 | 0.064 |
| 2 | 0.528 | 0.041* | -0.026* |  |  | 3.86 | 2,16 | 0.043 |

\* individual slope coefficients significant at $p < 0.05$
† total size of clan, including subadults.

The steepness of the female competition effect is evident from Fig. 4b, which plots the number of cubs per adult female controlling for number of males. A cubic equation gives the best fit (r^2^=0.635, F_3,15_=8.70, p=0.001 2-tailed). Notice the steepness of the initial decline, indicating a very rapid reduction in fertility across the range 1-5 females; from around five adult females, fertility enters a distinctive asymptotic phase, before returning to a steep linear decline again once there are more than ∼20 females in the clan. Judging by Fig. 4a, a residual fertility of -0.5 is probably equivalent to the demographic replacement rate, so that this plateau represents the minimum for survival as a clan. The plateau implies that hyaena females are able to exploit some kind of behavioural strategy that buffers them against the full force of the infertility trap, but that this solution does not protect them once there are more than about 20 females in the clan. Once clan size exceeds 20 females, the only way out of the infertility trap is probably clan fission. This then suggests that the clan enters an oscillatory phase in which it cycles through a nonlinear oscillator in which slow growth is followed by precipitate fission once the clan is large enough to yield two daughter clans with an average size of ∼10 females.

To test whether it is female alliances that help reduce the effects of the infertility trap (as they do in primates), I plotted lifetime fertility (cumulative number of cubs) derived from pedigrees for 31 adult females belonging to one large clan (the Talek clan in the Serengeti Park, Kenya) that has been studied over a 29-year period (Holecamp et al. 2012). Data are limited to the five original matriarchs and their daughters, all of whom could in principle be mothers or grandmothers (and in some cases, great-grandmothers). Fig. 4c plots completed family size (as of the date of the final census) against individual females’ within-matriline dominance rank. The horizontal line demarcates the effective replacement rate (two cubs over a lifetime, adjusted by the observed average cub survival to 18 months of 75%: Hofer & East 2003). Females whose lifetime output falls below this value (2.67 cubs) will not contribute enough surviving cubs to the next generation to replace themselves. The overall regression line is highly significant (r^2^=0.632, t_29_=-7.05, p<0.0001 1-tailed). The regressions for the individual matrilines are all negative, though only four are individually significant (p≤0.005, with p=0.125 for the exception, 1-tailed tests). That the data for the matrilines overlap rather than forming a linear sequence based on overall rank indicates that matrilineal alliances go some considerable way to buffering their members against the stresses of living in super-large groups with many females. However, this protection is not perfect, as is indicated by the fact that most of the data for the dominant matriline (filled circles) lie above the overall regression line, whereas those for the lowest ranked matriline (unfilled squares) mainly lie below the line, with the three mid-ranking matrilines in between.

The regression line in Fig. 4c intersects the replacement rate at a cohort size of 4.4 adult females (i.e. a value very close to that implied by Fig. 4b). In effect, groups with more than 4-5 females will not breed at rates high enough to maintain clan size at demographic stationarity (i.e. the clan will decline in size). If we take 4.4 as the limiting number of breeding females and add to this the number of adult males, subadults and cubs that we would expect to see in a clan with this number of adult females based on the demographic data for the sampled clans (Fig. S1), we get a total clan size of 12.2 (4.4 adult females, 2.4 adult males, 3.2 subadults and 2.2 cubs). Note how close this is to the commonest clan size in Figs. 1 and 3. Holecamp et al. (2012) report a mean size of 4.2 (range 2-7) adult females for five matrilines in the Talek clan. In effect, this represents the natural group size for the species. Larger clans can be engineered when needed, but, as in the primate case, require behavioural mechanisms to reduce stress levels – or unusually rich foraging conditions that will increase birth rates so as to offset the losses incurred by the infertility trap.

In the kinds of habitats occupied by spotted hyaena, we would expect clans to be larger in richer habitats (i.e. those with higher densities of prey), and these will typically be wetter and cooler. I test this as directional hypotheses in a multiple regression. Large groups are indeed found in higher rainfall habitats (slope = 0.034±0.017, t_18_=2.00, p=0.031 1-tailed), but the predicted effect of temperature is weak and only found in East African populations (overall slope = 0.125, t_18_=0.08, p=0.971 1-tailed) (Fig. 5). In hyaena habitats, high rainfall is usually associated with woodland, and in Africa woodland habitats typically have higher prey densities than drier grassland habitats.

**Fig. 5.**
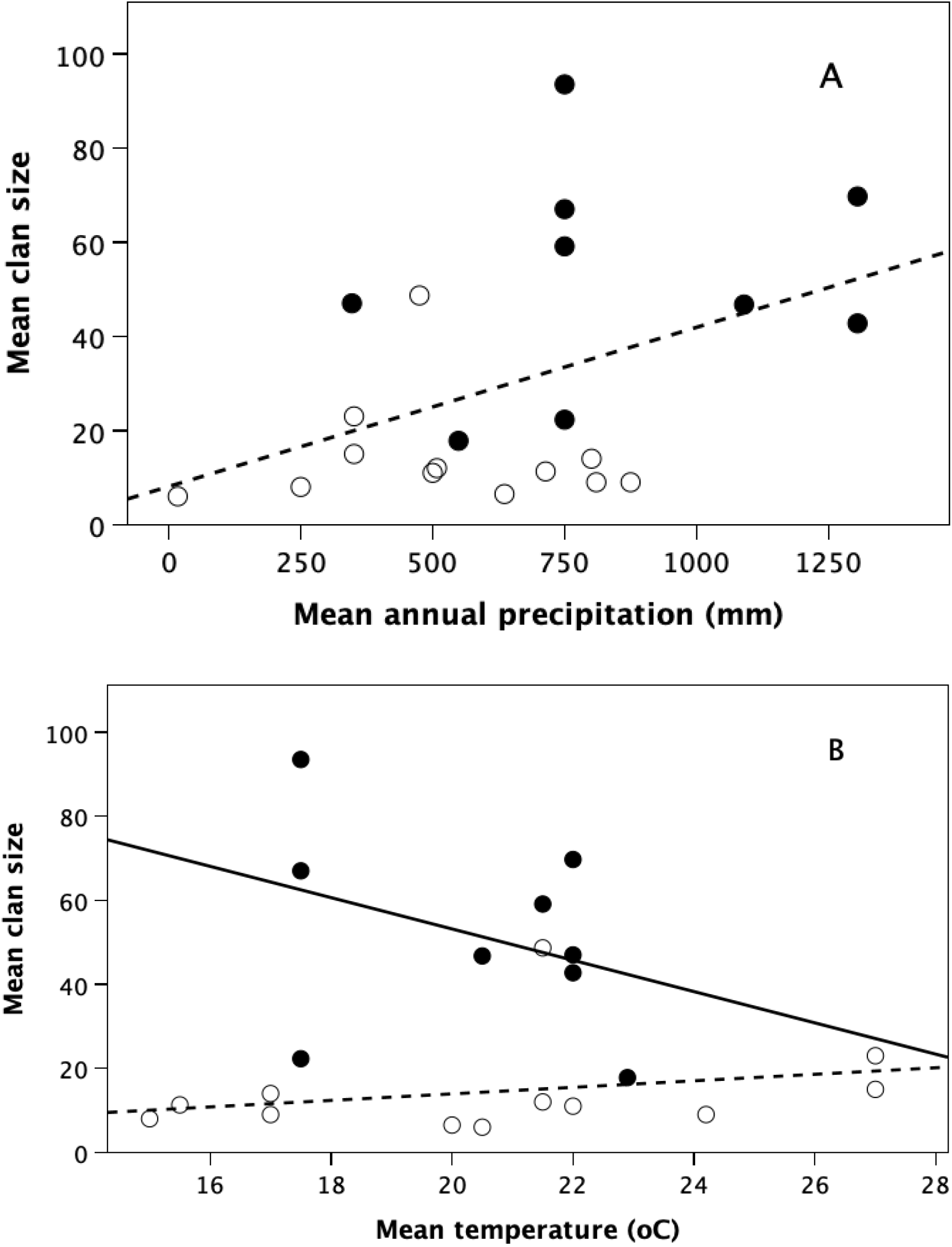
Mean clan size for East African populations (filled symbols) and southern African populations (unfilled symbols), plotted against (a) mean annual precipitation (mm) and (b) mean annual temperature (°C) of habitat. For precipitation, r^2^=0.183, t_18_=2.062, p=0.053 (though this might be better described by a step change at around 750 mm annual precipitation); temperature has no independent overall effect (t_18_=0.076, p=0.941), though the East African sites on their own have a near significant negative effect that converges on the southern African values when mean temperature exceeds 27°C.

Note that single matriline clans seem to be more common in southern Africa (where habitats tend to be drier), whereas clan sizes are more variable in eastern Africa (where larger clans are associated with wetter habitats) (Fig. 6). Males are not themselves members of the females matrilineal alliances, but seem rather to map themselves onto the number of reproductive females present in what seems to be an ideal free distribution. More importantly, note that the very large clan sizes in eastern Africa are mainly due to the presence of large numbers of adult males in addition to many females. While southern and eastern populations do not differ in the number of females, the number of males in southern clans seems to reach an asymptotic value at ∼7.5 males (Fig. 6). This implies a shortage of males in the southern part of the species’ range, perhaps due to higher male mortality rates.

**Fig. 6.**
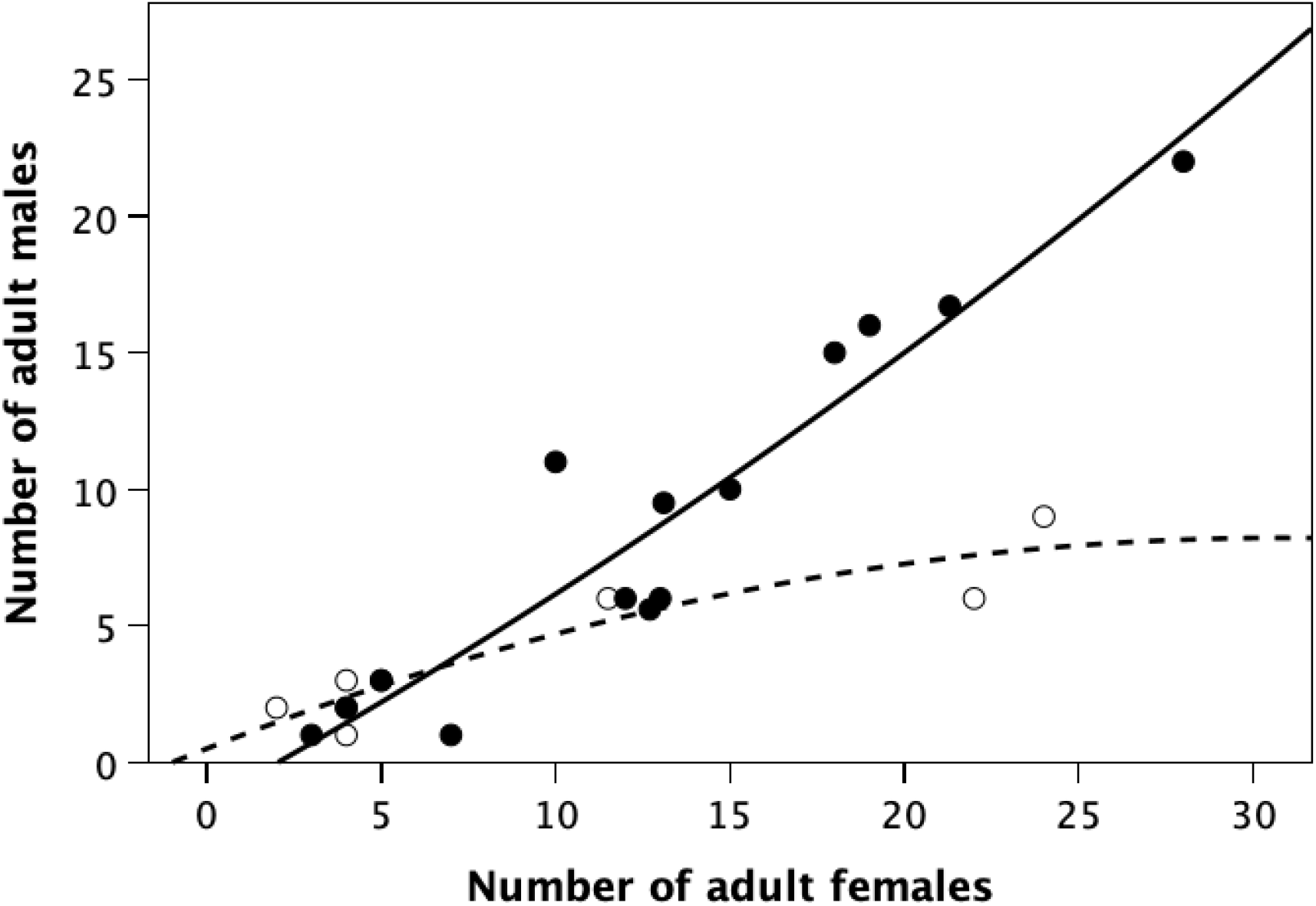
Number of adult males plotted against number of adult females for individual clans in East African (filled symbols) and southern African (unfilled symbols) sites. Quadratic regression: East African sites, r^2^=0.894, F_2,11_=46.25, p<0.001; southern African sites, r^2^=0.860, F_2,5_=15.40, p=0.007. The quadratic relationship for the East African sites does not differ from a linear relationship.

## Discussion

The distribution of group sizes in freeliving spotted hyaena populations is multimodal, and best described as forming two, perhaps three, overlapping distributions rather than a unimodal normal (or Poisson) distribution. A multimodal distribution is indicative of a multilevel social system. The lower cluster, with clan sizes in the region of 10-15 animals, is not only the most common numerically but seem to be more stable than clans of larger size. The distinctive phase shift in the slope parameter at a clan size of 15 (Fig. 3) sets an upper limit on the size of this lower cluster. The size of this lower cluster seems to be the outcome of an infertility trap that places a limit on the number of females who can live together as a group. The infertility trap is linear rather than ∩-shaped as in anthropoid primates (Dunbar & Shultz 2021b), with a limit at ∼4.5 females (equating to a total clan size of 12-15).

When a clan exceeds this size, it must either undergo fission or, if it is to continue increasing in size, find ways to mitigate the stresses that underpin the infertility trap. That clans do undergo fission is confirmed, in different populations, by Holecamp et al. (1993) and Höner et al (2005). In the three observed cases of fission, it was largely young, low-ranking females that emigrated (i.e. those likely to be doing less well reproductively) from large clans (14, 16 and 22 adult females, respectively). In some cases, they took juvenile offspring with them. Large clans (at least up to ∼150 in total size) are, however, evidently possible. In such cases, they become highly substructured into what are, in effect, matrilineal alliances (Holecamp et al. 1997, 2012). An analysis of association networks for one large clan revealed five matrilineal clans, plus three low-ranking female isolates. In effect, a clan size of 12-15 marks a point of transition between what we might consider the basal social unit (built around one matriline) and a multilevel unit consisting of several matrilineal sub-clans that can hunt semi-independently. This matrilineal structure allows females to form defensive alliances (Vullioud et al. 2019) not unlike those found in cercopithecine primates) that buffer their members against the infertility trap by preventing members of other matrilines from harassing them. Wahaj et al. (2001) report that spotted hyaenas exploit a number of behaviours like reconciliation that primates use to regulate their alliances and minimise the risk that alliances will break up as a result of aggression. How stable these very large clans are is not well known, and the fact that there are very few clans >90 members (and none at all with >150) suggests that fission is increasingly likely as clans approach this size.

While adult males seem to distribute themselves among the clans on an ideal free basis in response to mating opportunities (Höner et al. 2007), the South African clans had significantly fewer males per female than those from East Africa (Fig. 6). Whether this is because southern females are able to prevent new males from joining once they achieve numerical dominance (at about 15 females) or because high male mortality in these populations means that there are fewer males to distribute around the local clans is not clear. Watts & Holekamp (2008) provide evidence that competition with lions reduces clan growth rates, at least in East Africa (see also Höner et al. 2002). If lion densities are higher in the southern African habitats, and males are less risk-averse than females when attempting to scavenge from lion kills, it may well be that high male mortality is the explanation. Turner et al. (2020) have shown that more risk-averse hyaena are less likely to survive (although, in their rather specific experimental context based on between-clan conflict, they did not find a sex difference). More detailed empirical evidence is required to decide between these options, although, given the results in Table 1, the second might seem the more likely explanation.

Given the infertility trap, animals will only live in large social groups when there is a benefit from doing so. Table 1 indicates that having more males around does have a positive effect on fertility. This is evident from the contrast between the very large clans (total size >40) in southern Africa compared to East Africa: the former have much lower fertility than the latter (t_6.4_ = 8.44, p=0.0001). This might be because males help to provision mothers and cubs or because they help defend them against ecological competitors (neighbouring clans, or lion or African wild dog packs). Since males map themselves onto female groups on an ideal free basis (presumably for mating opportunities), females must be able to defer fission if they are to benefit by having extra males in the group. However, they may only be able to do this in habitats that are rich enough to support a denser prey population or prey species of larger body mass (Kruuk [1972], for example, found that the size of prey killed correlated with the number of hunters – in effect, clan size).

This suggests two predictions that can be tested: (1) the presence of more males allows subordinate females to reproduce more successfully if males help to provision them and (2) females hunt more often for themselves in small clans than in larger ones. Further issues which would benefit from additional empirical data are whether (a) males are more risk-prone in challenging other carnivores for their prey, (b) males are more likely to incur fatal injuries when hunting down large, dangerous prey, and (c) specific males favour provisioning particular females (or particular matrilines). The latter issue might suggest that matrilines, with attached males, form genuinely semi-independent sub-networks with large clans that both den and hunt separately at least part of the time. This would amount to exploiting the same kind of fission-fusion system found in gelada and hamadryas baboons (Dunbar & MacCurran 2019), and would explain why the fertility rates for different matrilines overlap in the way they do in Fig. 4c.

The finding that spotted hyaena are susceptible to the infertility trap and exploit primate-like alliance strategies raises the possibility that other mammalian taxa might behave in similar ways. We know that equids, lions and meerkats all exhibit the same kind of ∩-shaped fertility curves as anthropoid primates (Dunbar & Shultz 2021b), suggesting that they too use female alliances to defer the infertility trap. However, these species resemble anthropoid primate genera that live in small/medium sized bonded social groups. The spotted hyaena bear a closer resemblance to those anthropoid primates like gelada and hamadryas baboons that live in very large fission-fusion groupings and rely on substructuring into semi-independent matrilineal alliances to manage the infertility trap. This strategy may be more common in other non-primate mammalian orders where relationships between individual animals are more narrowly focussed within an extended social group.

## Supporting information

Supplementary Results

