## Supplementary Results for "Optimal Social Group Size in Spotted Hyaenas (*Crocuta crocuta*): Insights into a Multilevel Society"

**R.I.M. Dunbar**

**Supplementary Information**

Figure S1 plots relationships used to calculate demographic data for the characteristic composition of limiting clan size in the small cluster identified by Figs. 1 and 3. Regression equations for these are given in Table S1.

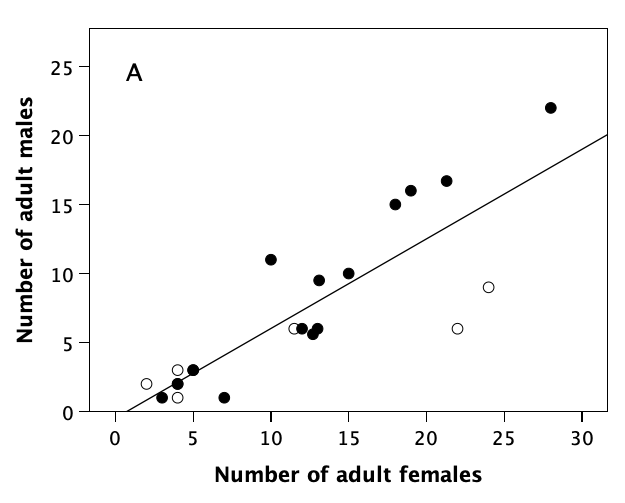

**Figure S1**

Number of (a) adult males, (b) subadults and (c) cubs in individual clans from East Africa (filled symbols) and southern Africa (unfilled symbols). Regression line is the overall OLS regression. Source of data: Supplementary Datafile 1.

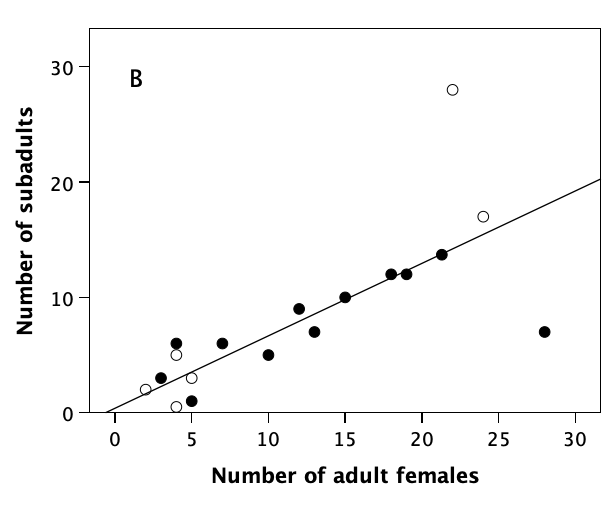

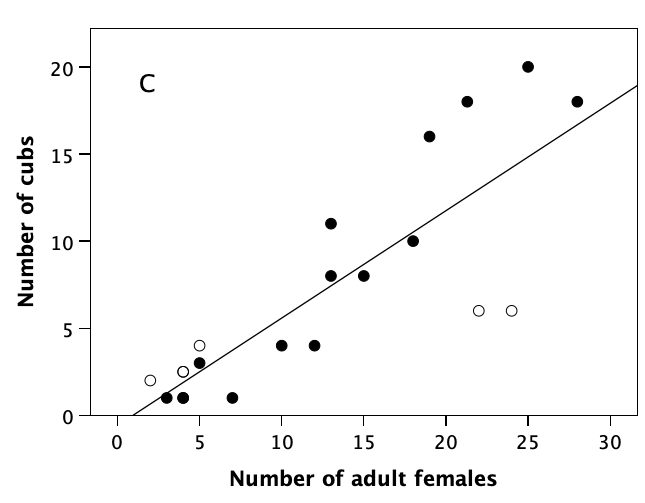

**Table S1**

Regression analyses for relationship between different demographic cohorts in a clan and the number of adult (i.e. breeding) females, for the distributions in Figure S1.

Dependent

variable	Equation	r^2^	t	df	p

Adult males	-0.471 + 0.649*Fems	0.713	7.04	18	<0.001

Subadults	0.385 + 0.618*Fems	0.571	4.76	16	<0.001

Cubs	-0.581 + 0.616*Fems	0.694	6.57	18	<0.001

Fems = number of adult females in clan
